# Larp1 supports brain growth and spatial memory via post-transcriptional control of the translation machinery

**DOI:** 10.1101/2025.10.09.681478

**Authors:** Mark S. Williams, Maegan J. Watson, Carson C. Thoreen

## Abstract

In the brain, tight regulation of the translation of mRNAs is essential for development and plasticity. The translation machinery itself is largely encoded by mRNAs with terminal oligopyrimidine (TOP) motifs, which can be post-transcriptionally controlled by the mTOR signaling pathway. In neurons, these mRNAs are selectively enriched in axons, dendrites and synapses, suggesting local functions for their regulation. Here, we use a brain-specific knockout of the mTOR effector and TOP mRNA binding protein, Larp1, to uncover its role in brain development and behavior. Loss of Larp1 significantly decreases brain mass and reduces the density of neurons. We find that TOP mRNAs levels are depleted by more than 50% and selectively lost from synapses, reversing the enrichment that occurs when Larp1 is present. In behavior tests, Larp1-deficient mice are severely impaired in spatial learning and memory. These results demonstrate a critical role for Larp1 in maintaining the levels of essential mRNAs necessary for brain growth and highlight the importance of post-transcriptional regulation by mTOR for normal learning and memory.

## Main text

The translation machinery is a complex system of ribosomes and translation factors that perform the essential task of synthesizing protein from mRNA. Aside from its fundamental function in gene expression, global regulation of the translation machinery can trigger context-dependent changes in cellular biology. In the brain, changes in levels of the translation machinery direct cell fate trajectories and brain development (1). In sub-cellular compartments, local control over levels of protein synthesis is involved in the growth of axons and dendrites as well as synapse development and plasticity (2). How levels of the translation machinery are adapted in the brain across different developmental stages, cell types, and sub-cellular compartments remains uncertain.

A conserved system for regulating the synthesis of the translation machinery is through the post-transcriptional regulation of mRNAs with terminal oligopyrimidine (TOP) motifs (3). TOP-containing mRNAs (TOP mRNAs) encode ∼200-300 proteins that include essentially all ribosomal proteins, key translation factors, and some nuclear-encoded mitochondrial proteins (4). The motif itself is a short pyrimidine-rich sequence at the 5′ termini of mRNAs, adjacent to the cap structure (3). It invariably begins with a +1 C followed by a series of 4-14 pyrimidine nucleotides (3). When appended with TOP motifs, the translation and stability of mRNAs becomes dependent on the mTOR Complex 1 (mTORC1) pathway, a sensor of nutrients and other growth signals that is linked to neurodevelopmental disorders (5). The result is that these mRNAs are translated normally when mTORC1 is active but are translationally repressed and stabilized when mTORC1 is inhibited. The regulation of TOP mRNA translation and stability are both mediated through an RNA-binding protein called La-related Protein 1 (LARP1) (6–8). When mTORC1 is inhibited, Larp1 binds the 5′ end of TOP mRNAs and sequesters them from the translation machinery by preventing access to the 5′ cap (7, 8). This represses the translation of these mRNAs, while also stabilizing them by preventing deadenylation through interactions with the poly-A tail (9, 10). This system allows cells to rapidly and locally control the synthesis of the translation machinery through a unified mechanism.

While TOP mRNAs are broadly expressed, they are particularly enriched in distal projections, including axons, dendrites and at synapses (11–18). In cultured neurons, where neurites can be separated from cell soma, the enrichment of TOP mRNAs in neurites is at least partly dependent on Larp1 (13, 15). Despite the ubiquity of TOP mRNA localization to distal regions of neurons, the biological significance of this enrichment remains uncertain. One hypothesis is localized TOP mRNAs are important for maintaining translation capacity at remote locations.

Although mature ribosomes are assembled in the nucleus, some ribosomal proteins on the surface of ribosomes can be exchanged at these distant sites, allowing damaged ribosomes to be repaired with locally produced replacement proteins, extending their lifetime and maintaining translation capacity (12, 17). Other TOP-encoded translation factors don’t require nuclear assembly and could contribute to the maintenance of the translation machinery or mitochondria at distal locations independent of ribosomes (4). Maintaining remote translation capacity may be important for the growth or remodeling of distal sites.

Brain disorders associated with mTORC1 hyper-activation offer additional insights into possible functions of TOP regulation. Under these conditions, Larp1 is inactivated and TOP mRNAs are constitutively translated (19). The prototypical example of these disorders is tuberous sclerosis complex (TSC), which is linked to heterozygous mutations in the upstream mTORC1 repressors TSC1 or TSC2 (20). In the human brain, TSC is characterized by the formation of focal lesions of dysmorphic cells, including giant neurons that have likely lost the remaining functional TSC1/2 allele. Affected individuals suffer from severe seizures, cognitive disability, and a high incidence of psychiatric disorders (20). In mice, deletion of Tsc1 in neurons leads to severe seizures and enlarged and dysplastic cells throughout the brain (21). Hypomorphic mutations in Tsc2 also increase in neuronal soma size and lead to learning deficits (22). Tsc2^+/-^ mice similarly show learning deficits (23) along with increased translation of TOP mRNAs (24). These observations suggest that the constitutive translation of TOP mRNAs, through inactivation of Larp1, might support neuron overgrowth and altered behavior.

In this study, we sought to determine the function of TOP mRNA regulation in the brain using a conditional mouse model with brain-specific deletion of Larp1 (Larp1 cKO). Our expectation had been that Larp1 inactivation would amplify brain and/or neuron growth by increasing TOP mRNA translation downstream of mTORC1, similarly to disease mutations that hyper-activate mTORC1 (20). In contrast, we find that the brain mass of Larp1 cKO mice is significantly reduced. TOP mRNA translation is increased in Larp1 cKO brains, but TOP mRNA levels are strikingly diminished by more than 50%. This depletion is amplified at synapses, where depletion of some TOP mRNAs exceeds 80%, and abolishes the enrichment observed in normal neurons. In behavioral tests, Larp1 cKO mice are markedly impaired in spatial memory. Overall, these results support the hypothesis that a central function of Larp1 is to sustain levels of TOP mRNAs, particularly in subcellular compartments distant from cell bodies of neurons, to support brain growth and plasticity.

## Results

### Loss of Larp1 in the developing brain reduces brain and body mass

To begin, we crossed mice expressing the cre recombinase from the Nestin promoter (Nestin-cre) with mice carrying a floxed allele of Larp1. Nestin-Cre is expressed in early (∼E9-11) progenitors of neurons and some glia, leading to widespread recombination of floxed alleles throughout the central and peripheral nervous system (25). Knockout mice (Larp1 cKO) are viable and born at the expected Mendelian ratios (Fig 1A). Larp1 protein levels are reduced by ∼50% throughout the brain of cHet mice and fully depleted in Larp1 cKO mice (Fig 1B). Body weights of Larp1 cHet mice are similar to WT controls, indicating that Larp1 protein levels are not normally limiting for overall growth. In contrast, the body weights of Larp1 cKO mice are significantly reduced (∼10%, p < 0.005) (Fig 1C). Brain weights of Larp1 cKO mice are also reduced (Fig 1D). The brain weights of cHet mice are also slightly reduced, although this difference reaches the level of significance only in male mice (Fig 1D). Brain-to-body mass ratios are decreased in Larp1 cKO mice, indicating that the loss of Larp1 has a primary effect on brain growth (Fig 1E). Nonetheless, overall cytoarchitecture was preserved in Larp1 cHet and cKO brains (Fig 1F). Overall, these results show that the loss of Larp1 significantly reduces brain growth without major disruptions in cytoarchitecture.

**Fig 1.**
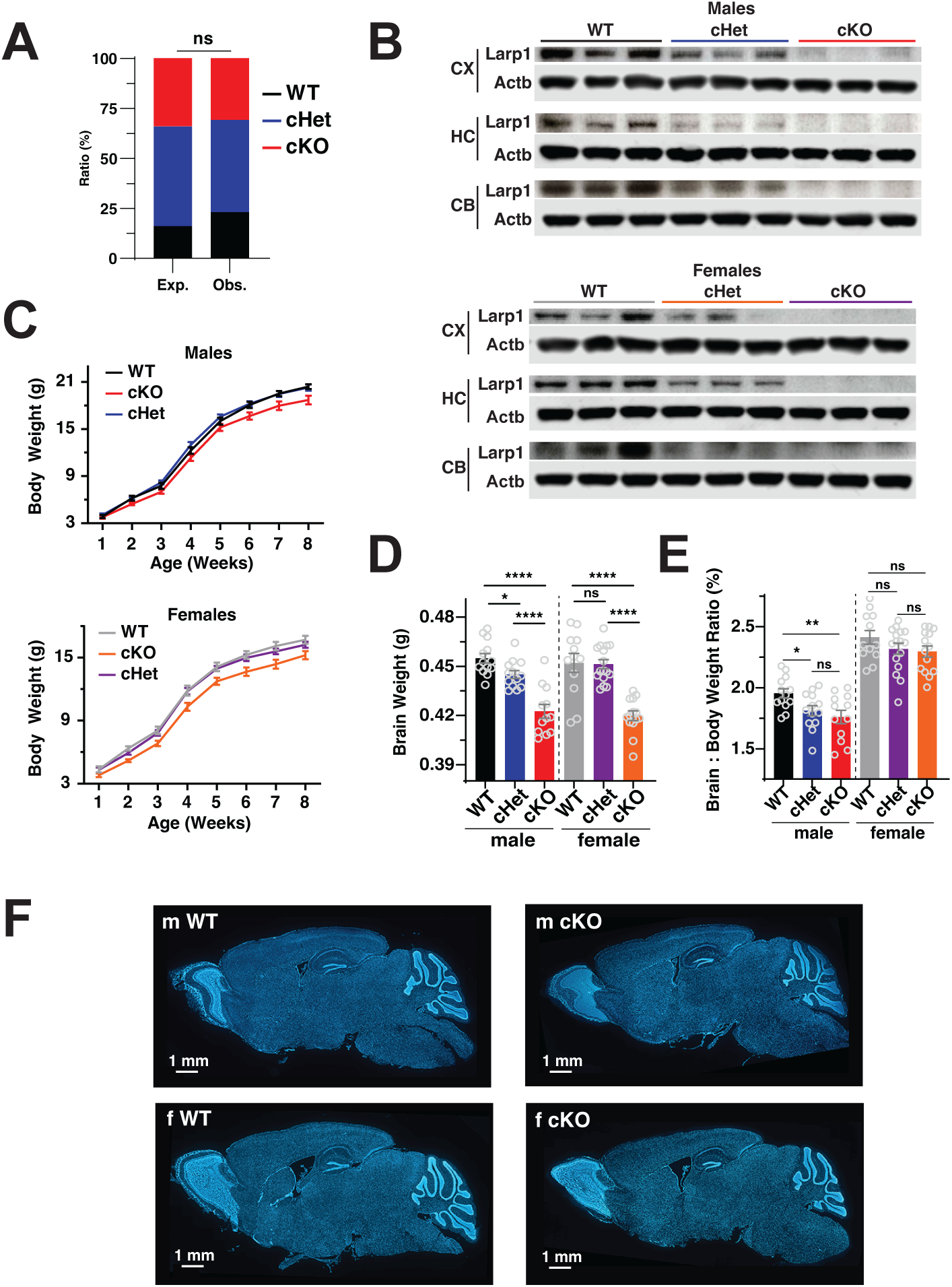
Conditional deletion of Larp1 in the mouse brain reduces brain and body size. **A.** Larp1 cKO mice are born at the expected ratios. Expected and observed frequencies of the indicated genotypes are shown (12 WT, 24 cHet and 16 cKO). Significance by Fisher’s exact test. **B.** Levels of Larp1 are reduced in brains of Larp1 cHet and cKO mice. Levels of Larp1 in the cortex (CX), hippocampus (HC) and cerebellum (CB) of male and female mice with the indicated genotypes were analyzed by western blotting for the indicated proteins (n = 3 for each genotype). **C.** Loss of Larp1 reduces body weights of mice. Mean weights of male and female Larp1 WT, cHet and cKO littermate mice at the indicated ages. Error bars are SEM. Significance determined by repeated measure one-way ANOVA with Tukey’s multiple comparison test correction. m WT vs m cKO p = 0.0027, m cHet vs m cKO p = 0.0005; f WT vs f cKO p < 0.0001, f cHet vs f cKO p < 0.0001. **D.** Brain mass is reduced in Larp1 cKO mice. Weights of brains from PBS-perfused 2-4 **mo** mice of the indicated genotypes (data are mean +/- SEM). For males, n WT = 13, n cKO = 12, n cHet = 13. For females, N WT = 12, N cKO = 14, N cHet = 16. Significance by unpaired t-tests, *p <0.05, **p<0.01, ****p<0.0001. **E.** Brain mass is disproportionally reduced in Larp1 cKO mice. Brain weights from (D) were normalized to body weights for the indicated genotypes. Significance by unpaired t-test, *p <0.05, **p<0.01, ****p<0.0001. **F.** Representative DAPI-stained sections of brains from the indicated genotypes.

### The density of neurons is reduced in Larp1 KO brains

The reduced brain weight of Larp1 KO mice could reflect a reduction in either the size or number of neurons. To quantify these properties, we used CellPose to analyze images of the isocortex, caudate putamen, and hippocampus from male WT and cKO mice (Fig 2A) (26). As expected, layers 2-3 of the isocortex are primarily populated by large pyramidal neurons that are relatively sparse (Fig 2B and C). Layer 4 neurons are the smallest with the highest density.

**Fig 2.**
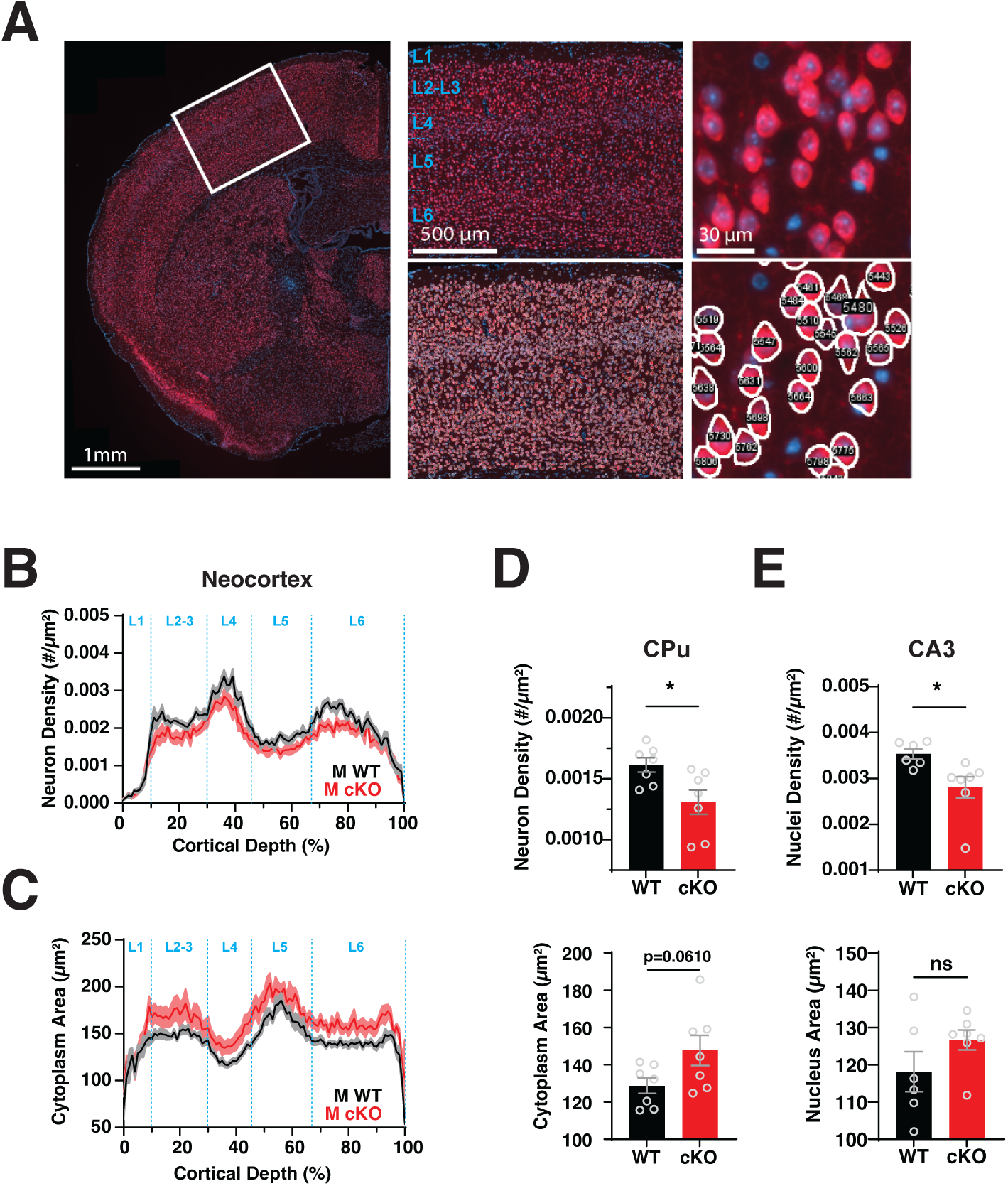
Reduced neuronal density and enlarged neuron size in Larp1 cKO brains. **A.** Overview of neuronal size and density analysis. Left panel: example WT 10 µm coronal section stained for neurons (NeuN) and nuclei (DAPI). Region selected for analysis is indicated. Center panel: Example region selected for analysis before (top) and after (bottom) computational segmentation of NeuN+ cells. Boundaries of cortical layers are indicated on the upper panel. Right panel: Magnified regions showing examples of identified NeuN+ cells. **B.** The density of cortical neurons is decreased in Larp1 cKO male mice. NeuN+ cells from regions of the primary somatosensory region from male Larp1 WT and cKO mice were segmented and binned by position along the vertical axis in 1% increments. Plot shows mean number and SEM within each bin. Significance by REML group, p value = 0.024, n=7 WT and 7 cKO. **C.** Neuronal soma are enlarged in Larp1 cKO mice. Areas of NeuN+ cells in segmented regions from (B) for male Larp1 WT and cKO mice. Traces are means +/- SEM. Significance by REML group p value = 0.0257, n=7 WT and 7 cKO. **D.** Neuronal density is reduced in the caudate putamen of male Larp1 cKO mice. NeuN+ cells in the caudate putamen (CPu) of male Larp1 WT and cKO mice were segmented and analyzed as in (B). Top panel: density of NeuN+ cells from the indicated genotypes. Significance by t-test, n=7 WT and 7 cKO. Bottom panel: area of NeuN+ cells from the indicated genotypes. Significance by t-test, n=7 WT and 7 cKO. **E.** Density of nuclei is reduced in the CA3 region of the hippocampus of male Larp1 cKO mice. DAPI+ nuclei from the CA3 region were segmented and analyzed as in (B). Top panel: density of DAPI+ nuclei in the indicated genotypes. Significance by t-test, n=6 WT and 7 cKO. Bottom panel: area of DAPI+ nuclei from the indicated genotypes. Significance by t-test, n=6 WT and 7 cKO.

Layer 5 contained the largest pyramidal neurons with a corresponding low density. And layer 6 contains slightly smaller neurons with a slightly higher density than layers 2-3. This typical laminar organization is preserved in Larp1 cKO brains (Fig 2B and C). We see no evidence that the sizes of neuronal soma are decreased. Instead, the sizes of neuronal somas are increased by ∼12% (p < 0.05) across cortical layers (Fig 2B and C). Conversely, soma density is reduced by ∼12% (p < 0.05). A similar pattern is evident in the caudate putamen (CPu) (Fig 2D). In the hippocampus, the density of neurons is too high to computationally segment. However, the size and the density of nuclei (which correlates strongly with soma size across all regions) in the CA3 region follows a similar pattern, although the difference in size did not reach the level of significance (Fig 2E) (27). We also observed no decrease in the size of neuronal soma in female brains (Supplemental Fig 1). These results argue that the decreased size of Larp1 cKO brains cannot be explained by the decreased growth of neuronal somas. Instead, it likely reflects a decrease in the total number of neurons produced.

### The translation of TOP mRNAs is increased in Larp1 cKO brains

Because Larp1 is a translation repressor, its loss is expected to increase TOP mRNA translation (4, 6, 8). To measure TOP mRNA translation in Larp1 cKO brains, we analyzed tissue from the posterior cortex using polysome profiling, which uses sucrose gradient fractionation to separate mRNAs bound by increasing numbers of ribosomes. Global polysome profiles show a slight increase in the abundance of single (80S) ribosomes in Larp1 cKO brains but are otherwise similar to profiles from WT cortex (Fig 3A). Normally, translationally-repressed Larp1-TOP mRNA complexes accumulate in 80S fractions of polysome profiles (28, 29). As expected, canonical TOP mRNAs (Eef2, Rps20, Rps6 and Rps25) are enriched in the 80S fractions in WT samples (Fig 3B). In Larp1 cKO cortex, TOP mRNAs are shifted to fractions with higher numbers of ribosomes, confirming a robust increase in their translation (4, 8). Given the increased translation of TOP mRNAs, we expected protein levels to also increase. Surprisingly, we find that levels of TOP-encoded proteins are not significantly higher in Larp1 cKO brains (Fig 3C). The increased translation of TOP mRNAs is thus paradoxically insufficient to significantly increase levels of TOP-encoded proteins.

**Fig 3.**
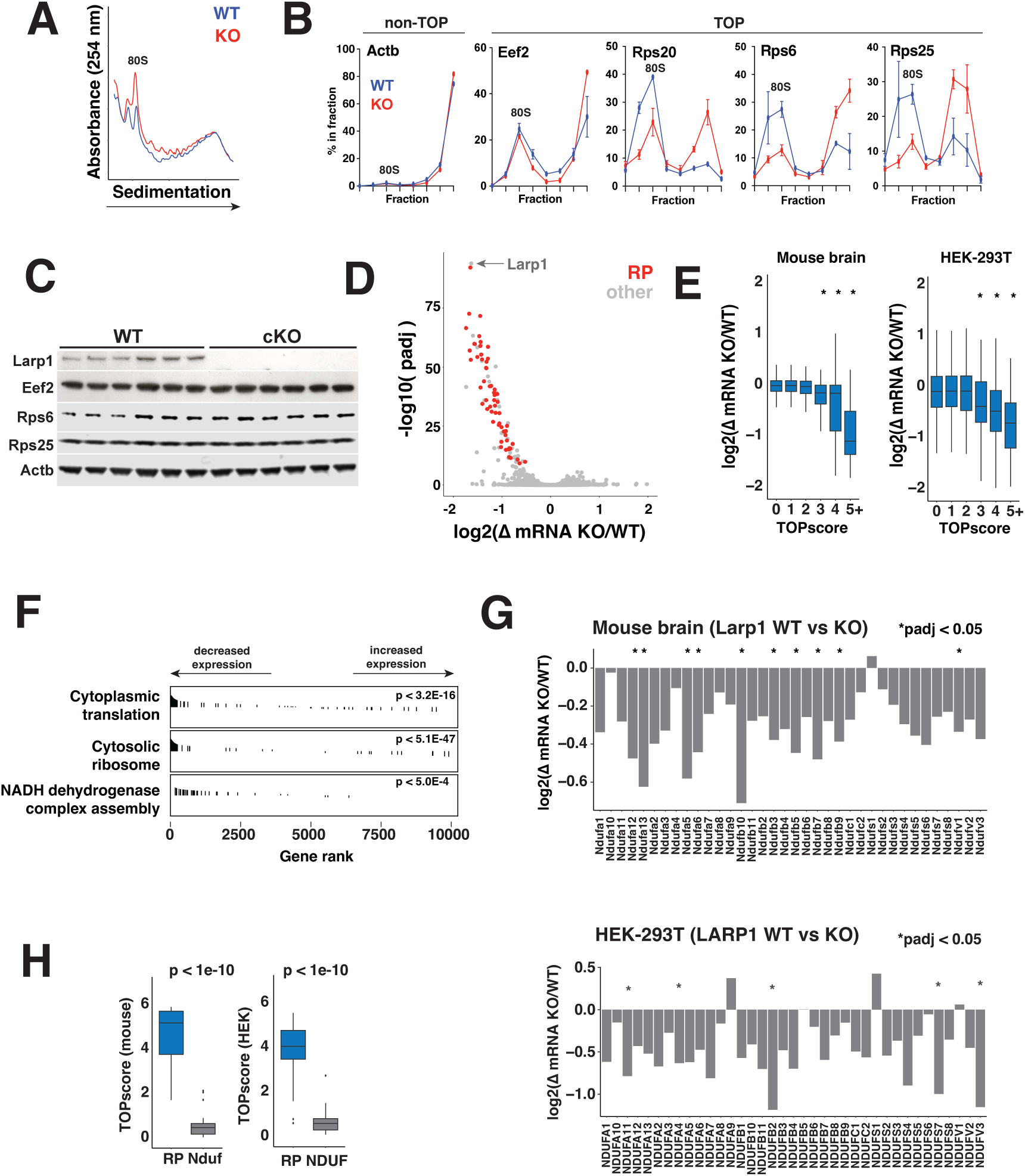
TOP mRNA translation and abundance are disrupted in Larp1 cKO brains. **A.** Polysome analysis of Larp1 WT and cKO cortex. Representative polysome traces for isocortex isolated from male Larp1 WT and cKO mice, homogenized and analyzed by sucrose gradient fractionation. **B.** The translation of TOP mRNAs is increased in Larp1 cKO brains. Levels of the indicated TOP and non-TOP mRNAs were quantified in fractions from (A) by qPCR**. C.** Levels of TOP-encoded proteins in the cortex of male Larp1 WT and cKO brains. Cortical samples from the indicated genotypes were analyzed by western blotting for the indicated proteins. (n=6 for each genotype). **D.** TOP mRNA levels are reduced in brains of Larp1 cKO mice. Similar posterior cortical regions from male Larp1 WT and cKO brains were analyzed by RNA-seq. Ribosomal protein (RP) mRNAs are highlighted in blue. n=3 for each genotype. **E.** Increased TOPscores are correlated with reduced levels in Larp1 cKO mice. Change in expression level between Larp1 WT and cKO brains and between LARP1 WT and KO HEK-294T cells for mRNAs with indicated TOPscores. TOPscores were calculated using FANTOM5 CAGE data for neonatal cortex and HEK-293 cells. Significance by ANOVA. **F.** Translation-related and mitochondrial mRNAs are depleted in Larp1 cKO mice. Changes in mRNA levels between Larp1 WT and cKO mice were analyzed by gene set enrichment analysis (GSEA). Significantly enriched gene ontology gene sets are shown. **G.** Complex 1 mRNAs are depleted in the absence of Larp1. Change in levels of the indicated Complex 1 mRNAs between Larp1 WT and cKO mouse cortex and between LARP1 WT and KO HEK-293T cells (n=3 for mouse cortical samples, n=2 for HEK-293T samples. * p_adj_ < 0.05). **H.** Distributions of TOPscores for ribosomal protein (RP) and Complex 1 NADH-ubiquinone oxidoreductase (NDUF) mRNAs in mouse cortical tissue and HEK-293 cells, as indicated. Significance by two-tailed t.test.

### Loss of Larp1 leads to depletion of ribosomal protein and mitochondrial protein mRNAs

In addition to repressing translation, Larp1 also stabilizes TOP mRNAs (9). We therefore wondered whether destabilization of TOP mRNAs in Larp1 cKO brains might be countering any increases in translation. To test this, we compared transcriptomes of posterior cortex from Larp1 WT and cKO mice. Overall, loss of Larp1 significantly altered only 293 mRNAs (padj < 0.05) out of 16,722 mRNAs detected (Fig 3D, Supplemental Table 1). Most (239) of these mRNAs are depleted and overwhelmingly composed of classical TOP mRNAs, including essentially all ribosomal proteins, key translation factors (e.g. Eif3 components, Eif4b, elongation factors), regulators of ribosome biogenesis (e.g. Nsa2), and mitochondrial proteins (e.g. Cox7a2l, Atp5j, Atp5e, Atp5h) (3, 4). We also found many depleted mRNAs that are not part of the canonical TOP family (e.g. Aimp1, Use1, 2410006H16Rik). However, closer inspection of transcription start site sequences for many of these mRNAs revealed clear TOP motifs (Supplemental Table 2, Supplemental Fig 2). Moreover, TOPscores, a metric that quantifies the strength of TOP motifs, are highly predictive of decreased transcript levels in Larp1 cKO (Fig 3E, Supplemental Table 2) (4). A similar effect is apparent in human LARP1 KO HEK-293T cells that we generated previously, underscoring the conservation of this function across cell types and species (Fig 3E) (8). Overall, the depletion of these and other TOP mRNAs is consistent with a major role for Larp1 in their stabilization through a mechanism that depends on the presence of TOP motifs.

We also find a significant number (93/239) of mRNAs that are reduced despite lacking clear TOP motifs (TOPscore < 3). This included additional mitochondrial protein mRNAs, including many components of Complex I (Fig 3F-I). While some mRNAs encoding mitochondrial proteins include clear TOP motifs, such as most components of the mitochondrial ATP synthase (Complex V), most encoding Complex I components do not, with the possible exceptions of Ndufb5, Ndufa6, and Ndufs2. Larp1 may stabilize these transcripts through recognition of other features, or their transcription may be indirectly repressed through other mechanisms that respond to decreases in other Larp1-target mRNAs. Even so, Complex I mRNAs are also depleted in LARP1 KO HEK-293T cells, arguing that Larp1 regulates their expression through a mechanism that is broadly conserved between mice and humans (Fig 3G).

### The loss of Larp1 reverses the synaptic enrichment of TOP mRNAs

In cultured neurons, the depletion of Larp1 reduces the enrichment of TOP mRNAs within neurites (13, 15). To test the impact of Larp1 on the synaptic enrichment of TOP mRNAs in vivo, we isolated synaptosomes from the isocortex of WT and Larp1 cKO brains. We confirmed that synaptosome preparations were enriched for the synaptic marker Psd95 (Fig 4A). As was observed in total cell extracts, synaptosome levels of TOP-encoded proteins were similar between Larp1 WT and cKO samples (Fig 4A). To quantify changes in synaptic transcriptomes, we analyzed synaptosome RNA using RNA-seq (Fig 4B). In WT animals, we detected 773 of 13993 coding mRNAs that were significantly enriched (padj < 0.05) more than 2-fold in synaptosome versus total mRNA libraries (Fig 4B). These mRNAs were overrepresented for gene ontology (GO) categories consistent with isolation of synaptic compartments (Fig 4C).

**Fig 4.**
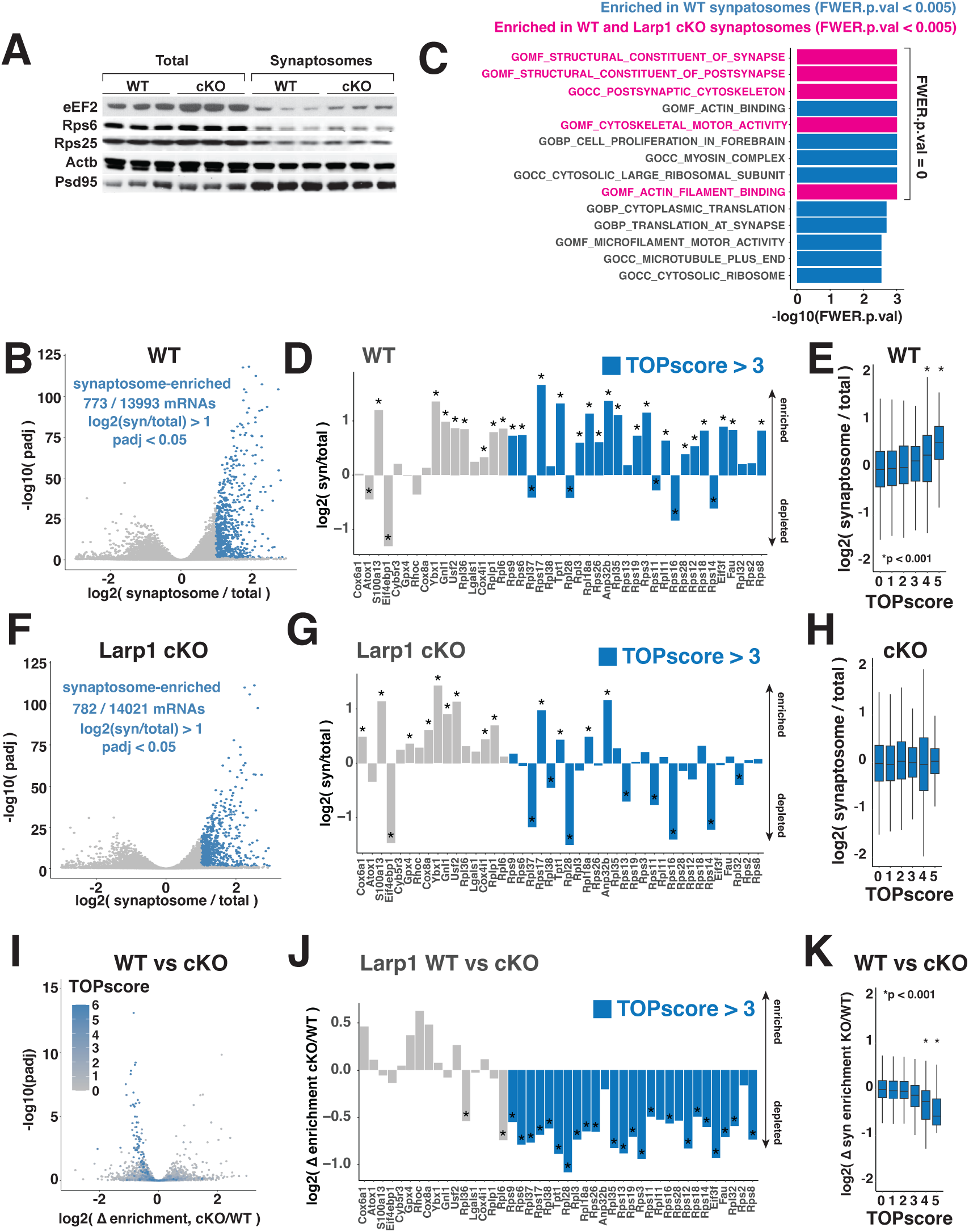
Larp1 inactivation depletes TOP mRNAs from synaptosomes. **A.** Levels of TOP-encoded proteins in synaptosomes. Total and synaptosome samples isolated from male Larp1 WT and cKO brains were analyzed by western blotting for the indicated proteins. **B.** TOP mRNAs are depleted in synaptosomes from WT brains. Levels of mRNAs in synaptosomes from WT brains were analyzed by RNA-seq and compared to total mRNA levels (n=3 for each genotype, total and synaptosome. Significance determined by DESeq2) **C.** Gene ontology categories overrepresented amongst synaptosome-enriched mRNAs. Synaptosome-enrichment values (synaptosome/total) were analyzed by gene set enrichment analysis (GSEA) for RNA-seq libraries from WT and Larp1 cKO mice. Significance levels were calculated using GSEA software. **D.** Enrichment of “core” neurite mRNAs detected in this dataset. Barplots show synaptosome levels of indicated mRNAs relative to total levels from analysis in (B). mRNAs are ordered by increasing TOPscore. Significance determined by DESeq2. **E.** Synaptosome enrichment correlates with TOPscore. Synaptosome enrichment for mRNAs determined in (B) was calculated for mRNAs with the indicated TOPscores. Significance determined by t-test between each class and mRNAs with TOPscores < 1. **F**. TOP mRNAs are depleted in synaptosomes from Larp1 cKO brains. Levels of mRNAs in synaptosomes from Larp1 cKO brains were analyzed as in (B). **G**. Enrichment of “core” neurite mRNAs in synaptosomes isolated from Larp1 cKO brains was analyzed as in (D). Synaptosome enrichment for mRNAs with the indicated TOPscores was determined as in (E). **H.** Change in synaptosome enrichment of mRNAs between WT and Larp1 cKO brains. Log2 fold-change and significance in synaptosome enrichment between genotypes was determined by DESeq2. **I.** Change in enrichment between WT and Larp1 cKO synaptosomes for “core” neurite mRNAs analyzed in (I). **K**. Change in synaptosome enrichment for mRNAs with the indicated TOPscores. Significances was determined by t-test between each class and mRNAs with TOPscores < 1.

Notably, GO classes related to ribosomes and translation were also significantly enriched, consistent with previous studies (13, 15, 18). We also detected enrichment of 26 of 43 mRNAs belonging to a “core neurite” transcriptome (18). This set contained 8 mRNAs without strong TOP motifs (TOPscores < 3), such as the calcium-binding protein S100a13 and the multi-functional protein Ybx1, and 18 TOP-containing (TOPscore > 3) mRNAs, most of which are classical TOP mRNAs that encode for ribosomal proteins or translation initiation factors (Fig 4D). Consistent with this observation, overall synaptosome enrichment was strikingly correlated with TOPscores (Fig 4E).

In Larp1 cKO synaptosomes, the overall number of coding mRNAs that are significantly enriched by more than 2-fold (782/14021, padj < 0.05) is similar to what we observed in WT synaptosomes (Fig 4F). These mRNAs remain enriched for GO categories related to synaptic function (Fig 4C). However, the enrichment of translation-related GO categories was lost. A similar pattern was evident in the “core neurite” transcriptome, where non-TOP mRNA enrichment was largely unchanged while TOP-containing mRNAs were remarkably depleted (Fig 4G). We note that Rpl36 and Rpl6 are depleted in Larp1 cKO synaptosomes despite having apparent TOPscores of less than 3 (1.81 and 2.77, respectively), but this likely reflects minor differences between the reference dataset used to calculate TOPscores and synaptosome mRNAs analyzed here. Globally, the correlation between enrichment and TOPscores was lost in Larp1 cKO synaptosomes (Fig 4H).

Overall, TOP motifs were a defining feature of mRNAs depleted in Larp1 cKO synaptosomes. 61/86 mRNAs that were significantly depleted from Larp1 cKO synaptosomes (padj < 0.05) had TOPscores greater than 3 (Fig 4I). TOP mRNAs amongst the “core neurite” transcriptome were similarly depleted in a Larp1-dependent manner (Fig 4J), while TOPscores were significantly associated with a Larp1-dependent decline in synaptosome enrichment (Fig 4K). The selective synaptic depletion of TOP mRNAs, but not other mRNAs whose total levels declined in Larp1 cKO brains (e.g. Nduf mRNAs), was confirmed by qPCR (Supplemental Fig 3A,B). These observations strongly support a model where Larp1 enriches ribosomal protein and other translation factor mRNAs at synapses through recognition of TOP motifs. We do find some examples (17/86) of mRNAs whose enrichment declines with the loss of Larp1 that appear to lack TOP motifs. The decreased synaptic enrichment of these mRNAs may reflect an indirect response to synaptic changes in Larp1 cKO brains. We also note that the enrichment of 19 mRNAs is significantly (padj < 0.05) increased in Larp1 cKO synaptosomes by more than 2-fold (e.g. Gpr6, Scn4b). These increases potentially reflect compensatory responses to Larp1-dependent changes in synaptic function. Nonetheless, the primary effect of Larp1 deletion on the synaptosome transcriptome is a significant and selective reduction in levels of TOP mRNAs that encode core components of the translation machinery.

### Spatial memory is impaired in Larp1 cKO mice

We next sought to determine how disruptions in TOP mRNA regulation impacted brain function by assessing behavioral differences in Larp1 cKO mice. Mouse models of mTORC1 hyper-activation, where Larp1 is expected to be repressed, have complex behavioral phenotypes, including altered anxiety, autism-like behaviors and impaired cognitive function (23, 30, 31). To assess the behavior of Larp1 cKO mice, we initially employed an open field test to measure exploratory drive and locomotive function (32). Larp1 cKO mice travel slightly but significantly less distance than control mice (Fig 5A). To assess whether the decreased distance travelled might reflect locomotor deficits, we measured fall latency using a Rotarod test with a steadily increasing rotation. The fall latency is not significantly different between Larp1 WT, cHET and cKO mice (Fig 5B). The loss of Larp1 therefore appears to slightly decrease exploratory behavior but does not appear to significantly impair locomotor function or learning.

**Fig 5.**
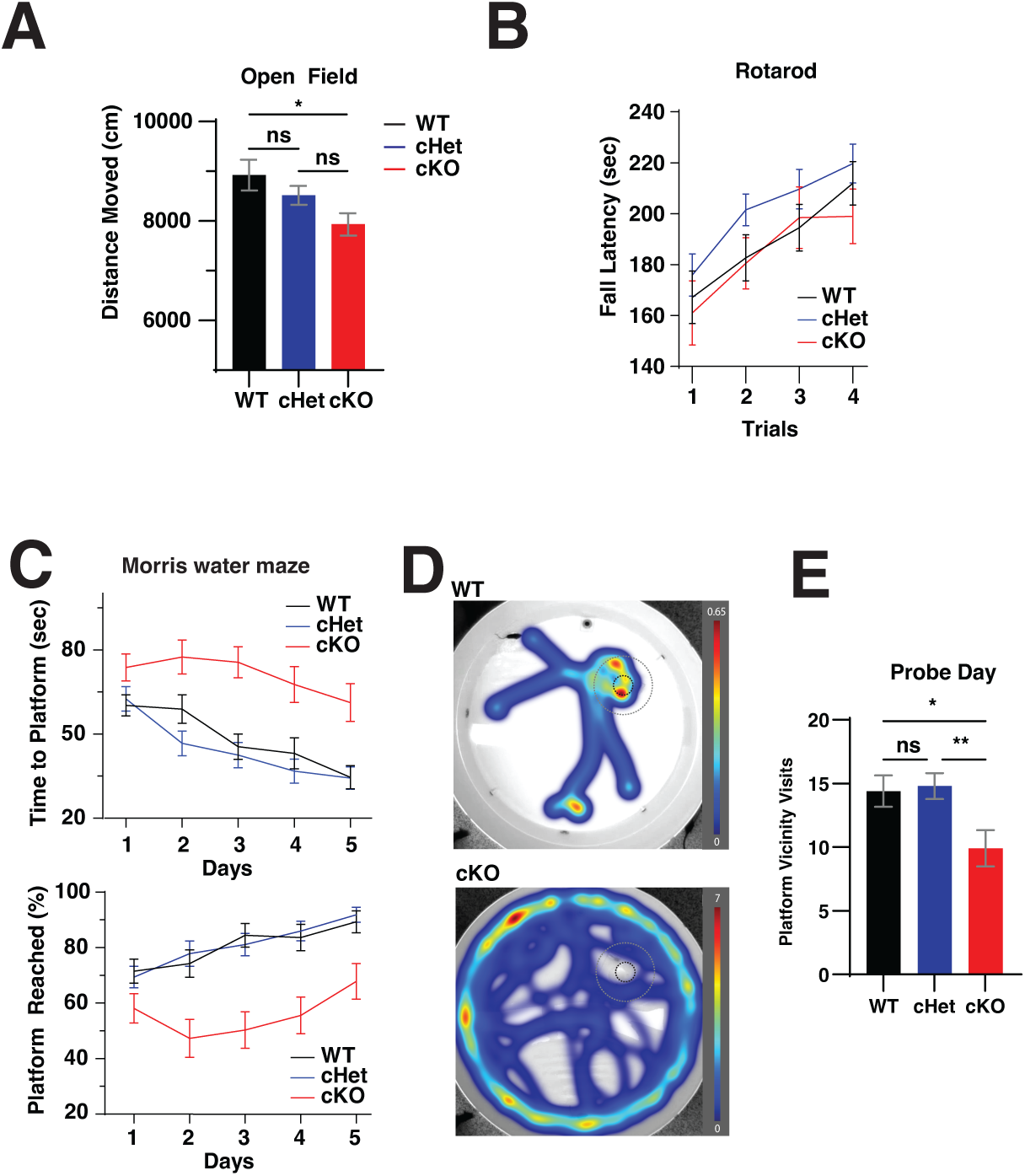
Behaviors of Larp1 cKO mice. **A.** Distance travelled for Larp1 WT, cHet, and cKO mice in an open field test. Distance traveled distance during the 30 min test time, binned by 5 min intervals. Significance by RM ANOVA (p = 0.0071). The male cHet group showed no significant difference with the cKO (p = 0.066) nor with the WT (p = 0.14). **B.** Locomotor learning is similar in Larp1 WT and cKO mice. Motor learning was assessed by rotarod test in ramp mode over 4 sessions from 5 to 40 rpm over 5 minutes. Data shown are mean +/- SEM for each session for the indicated genotypes. Significance by RM ANOVA. **C.** Increased escape latency for Larp1 cKO mice in a Morris water maze test. Top panel: Time to reach hidden platform for the indicated genotypes during 4 trials on each of 5 training days (n WT=45, n cHet=47, n cKO=34). Significance by RM ANOVA. Bottom panel: frequency of successfully reaching the hidden platform during the 2 min test. Significance by RM ANOVA. **D.** Representative examples of heatmaps for Day 5 trials. Heatmaps depict time at each position over 4 separate trials for a representative male Larp1 WT and cKO mouse. **E.** Visits to the platform vicinity (20 cm diameter) during a Day 6 probe trial for mice of the indicated genotypes. Significance by two-tailed t-test, n WT=45, n cHet=47, n cKO=34.

We next measured the performance of mice in a Morris water maze test, which is used to assess spatial memory (33). On each day of a 5-day training with 4 sessions per day, mice are placed in different quadrants of a 120 cm size pool and the time required to find the hidden (submerged) platform is recorded. As expected, control mice gradually improve over the course of the training, and by Day 5 reach a plateau in the time required to find the platform and reach 90% success rate. (Fig 5C). Larp1 cHet mice performed similarly to control mice (Fig 5D). In sharp contrast, Larp1 cKO mice take significantly longer to find the submerged platform and don’t have any more success at finding the platform at day 5 (Fig 5C). This deficit is evident even on the first day, which includes four separate trials. A representative trial from Day 5 shows how Larp1 cKO mice continue an essentially random search across the pool (Fig 5D). In a probe trial on Day 6, when the platform is removed, Larp1 cKO mice search less time in the vicinity of the original platform location than WT or cHet control mice (Fig 5E). Together, these results demonstrate that Larp1 cKO mice have significant deficits in spatial memory and learning.

## Discussion

In this study, we examined the function of TOP regulation in the mouse brain by inactivating Larp1, the TOP regulator. The global regulation of ribosomal protein and other TOP mRNAs has been implicated in brain development while their local regulation at distal locations has been proposed to support neuron growth and synaptic plasticity (34). Given Larp1’s function as a translation repressor of TOP mRNAs, we initially hypothesized that Larp1 loss would amplify growth phenotypes by increasing the translation of ribosomal protein and other TOP mRNAs similarly to mouse models of mTORC1 hyper-activation. Increased translation may still occur in specific contexts (for instance, in mature post-mitotic neurons) but the dominant molecular phenotype is a reduction in the global levels of TOP mRNAs that is exacerbated at synaptic sites, reversing the synaptic enrichment of these mRNAs. This reduction in TOP mRNA levels appears to be balanced by their increased translation, limiting the production of TOP-encoded proteins.

This global depletion of TOP mRNA levels offers a rationale for the low growth phenotype we observe, and also Larp1 loss-of-function phenotypes seen in flies, worms and plants (35–38). In each of these organisms, inactivation of Larp1 homologues significantly decreases growth. These slow growth phenotypes are reminiscent of other mutants with defects in ribosome biogenesis. For instance, ‘Minute’ Drosophila mutants, which are caused by mutations in ribosomal protein genes (39), are typified by slow development, reduced viability, low fertility, and short/thin bristles (40). Drosophila with homozygous dLarp mutations are also sterile and have similar short/thin bristle defects (35). In humans, mutations in ribosomal proteins and ribosome biogenesis factors lead to a family of disorders that are collectively known as “ribosomopathies” (41). Ribosomopathy phenotypes are variable, but mutations in RPL10 (42, 43), RPS23 (44), the ribosome biogenesis factor RBM28 (45), and the ribosome biogenesis factor AIRIM (46) all lead to microcephaly. This similarity to microcephaly we observe in Larp1 cKO mice suggests that Larp1 inactivation and subsequent depletion of ribosomal protein mRNAs might slow brain growth through similar mechanisms. The reduced growth potential of Larp1 loss-of-function mutants may also underly Larp1’s reported role as an oncogene in some tumor models (47).

The depletion of TOP mRNAs from distal subcellular compartments may also limit brain growth (18). While our analysis focused on synaptosomes, it seems likely that Larp1 also supports the enrichment of TOP mRNAs at axonal growth cones and in dendrites noted by previous studies (12, 13, 15–18, 48, 49). These results are in line with observations of Larp1 function in cultured neurons, supporting a model where Larp1 either directly transports TOP mRNAs to these distal locations or leads to their enrichment by preferentially stabilizing them (13, 15). Surprisingly, this depletion of TOP mRNAs from distal compartments does not appear to substantially impair the overall numbers of synapses, as levels of structural synaptic proteins (i.e. Psd95) are not globally altered (Fig 4A). However, growth defects may be limited in a subset of projects that are, for instance, beyond some threshold distance from cell nuclei. Larp1 is also required for mTORC1-dependent regulation of TOP mRNA translation and stability. TOP mRNAs might normally be translated only sporadically in response to transient mTORC1 activation. Such sporadic synthesis could be physiologically important without substantially alerting global levels of TOP-encoded proteins.

An intriguing question is whether the local deregulation of TOP mRNAs accounts for the impairment of spatial memory that we observe. At least some forms of synaptic plasticity depend on the local translation of mRNAs (34, 50–53). We wonder whether this requires the synaptic enrichment of TOP mRNAs. While the local translation of TOP mRNAs is unlikely to directly increase the local concentration of ribosomes, increased production of other TOP-encoded translation factors could enhance the capacity of the translation machinery by, for instance, increasing initiation rates. The regulation of TOP mRNAs with mitochondrial functions (e.g. Cox7a2l) may also be important for sustaining mitochondrial function at these distal sites, a process also linked to plasticity (54). Other mouse models of mTORC1 hyper-activation, where Larp1 is expected to be inactivated, show similar spatial memory deficits. For instance, Tsc2^+/-^ mice perform poorly in test of spatial memory (23). Similar to Larp1 cKO brains, the translation of TOP mRNAs is elevated in brains of these mice while TOP mRNA levels are reduced (24).

The translational control of TOP mRNAs is also lost in the absence of a second family of mTORC1-regulated translation repressors called 4E-binding proteins (4E-BPs), which are required to remove the eIF4F cap-binding complex and allow Larp1 to bind (8, 55, 56). As with Larp1 cKO and Tsc2^+/-^ mice, mice lacking 4E-BP2 are impaired in spatial memory (57). The inability to fully suppress the translation of TOP mRNAs in these models may therefore interfere with synaptic plasticity by locally altering levels of TOP-encoded translation factors near synaptic connections.

In summary, we find a major role for Larp1 in controlling the abundance and sub-cellular localization of TOP mRNAs in the brain, with important consequences for brain growth and function. The loss of Larp1 increases the translation of TOP mRNAs but also leads to striking reductions in TOP mRNA levels, particularly at synapses, limiting the capacity of neurons to produce ribosomes and other TOP-encoded translation factors and mitochondrial proteins. The resulting impairments in brain growth and spatial memory underscore the physiologic significance of TOP regulation and suggest that local regulation may be important for some types of synaptic plasticity. Many questions remain surrounding the molecular and cellular changes that underly these phenotypes. A key one is to sort out the specific contributions of changes in the translation machinery and mitochondrial function to the production and growth of neurons during development. A second is to understand how the depletion of TOP mRNAs from synapses and other distal sites impacts neuron growth and plasticity. Answering these questions will help to understand how local translation impacts neuron function and may also illuminate disease mechanisms in disorders such as TSC that can be targeted therapeutically.

## Methods

### Materials

Reagents were obtained from the following sources: TRIzol Reagent from Life Technologies; Protoscript II reverse transcriptase, NEBNext Ultra II Directional RNA Library prep kit, and NEBNext rRNA Depletion kit from New England Biolabs; DAPI from Cell Signaling Technology; iTaq Universal SYBR Green Supermix and Bradford Protein Assay from Bio-rad; and RNeasy Plus Mini Kit from Qiagen.

### Antibodies

**Table 1.** Antibodies used in this study.

| Target | Catalog # | Source | Concentration |
| --- | --- | --- | --- |
| NeuN | 94403S | Cell Signaling | IF 1:500 |
| Anti-mouse Alexa Fluor 647<br>Conjugate | 4410S | Cell Signaling | IF 1:500 |
| LARP1 | 14763S | Cell Signaling | WB 1:1000 |
| eEF2 | 2332S | Cell Signaling | WB 1:1000 |
| RPS25 | PA5-56865 | Invitrogen | WB 1:1000 |
| RPS6 | SC-74459 | Santa Cruz | WB 1:1000 |
| NDUFB10 | A3982 | ABClonal | WB 1:1000 |
| PSD95 | A0131 | ABClonal | WB 1:1000 |
| β-Actin | 3700S | Cell Signaling | WB 1:5000 |
| Goat Anti-mouse IgG 800CW | 926-32210 | LI-COR Bio | WB 1:5000 |
| Goat Anti-rabbit IgG 800CW | 926-32211 | LI-COR Bio | WB 1:5000 |
| Anti-Rabbit HRP Conjugate | 7074S | Cell Signaling | WB 1:5000 |
| DAPI | 4083S | Cell Signaling | 1 μg/mL |

### Transgenic mice

All animal experiments were approved and conducted in accordance with Yale Animal Resources Center and Institutional Animal Care and Use Committee (IACUC) regulations (protocol # 2019-20235). All mice (C57BL/6J) were housed in the Yale Animal Research Center of Yale University, in compliance with the US National Institutes of Health Guide for Care and Use of Laboratory Animals. Animals were housed with lighting on a 12h-12h light-dark cycle with 30–70% humidity and a 20–26 °C temperature. Animals had unrestricted access to water and standard rodent diet (ref #2018S, Teklad diets). Mice of either sex were used. Larp1 brain mutant mice were generated by crossing brain-specific Nestin-Cre mice acquired from The Jackson Laboratory with mice having loxP-flanked LARP1 gene (a gift from Ken Inoki) (58). Nestin-Cre mice have albinism and mice used for this study were selected from black furred animals of at least 2 generations down and heterozygous for Nestin-Cre. All genotypes were determined by PCR. Breeding pairs were cousins at the closest. Mice were weaned at 21 days postnatal and housed in groups of 2 or 3. All mice had unlimited access to food and water.

### Behavior exclusion criteria

Experimenter was blind to the mice genotype at the time of the experiment. Mice underwent handling no earlier than 7 weeks of age, and behavior no earlier than 8 weeks of age. No mice used for behavior was previously used for breeding. Mice had to be in good health and their cage free from obvious fighting to be included in behavior tests. No behavior test was run on the day of a cage change.

### Weights

Mice were weighed once a week, not more than 1 day prior or after their birth date. Weight was measured with a precision of 1 decimal. Animals used for weights always included at least 1 cKO mouse as littermate.

### Handling and habituation

Prior to handling or any behavior test, mice were brought in the same room and left to acclimate in the same conditions as the handling/ test with dim white lights and quietness, for at least an hour. Mice were nearly always found to be sleeping before the start of the procedure. No behavior test was conducted the same day as a cage change. Every day for 5 consecutive days, handling was performed by gently picking up the mice by the base of the tail and letting them explore both gloved hands, roughly 30 cm above their cage, for 2 minutes. The process of gently picking them up by the tail was repeated 3-4 times during those 2 minutes. All mice displayed a significant reduction in stress signs and no urination by day 3 of handling.

### Open Field

At the start, mice were placed at the center of the open field. Mice were free of movement and video recorded for 30 minutes. All videos, tracking and analysis was done with Ethovision XT (Noldus). Mice underwent the test only once. Mice of different sex were not run simultaneously. A maximum of 4 mice were run at the same time. Lighting was dim, white, and even across the entire 40cm x 40cm fields. Sides of the open field apparatus (Noldus) were 40cm high and center zone was defined as further than 8 cm of any wall. The test was conducted in a specific behavior room, quiet, with dark painted walls and dim overhead white light. The experimenter stayed more than 2 meters away while the test was running. The apparatus was cleaned with 70% ethanol and dried before every session.

### Rotarod

The day prior to the test, the mice underwent a training session in the same room and conditions. The training consisted of 2 sessions at constant speed of 5 rotations per minute (rpm), for a duration of 2 minutes and with an interval of 2 hours.

The test consisted of 4 sessions, at 30 minutes interval. At the start of each session, 1 to 5 mice were placed on the rod rotating at 5 rpm for 10 to 15 seconds before the start of each run. Runs consisted of an increase in rotation of the rod with a start at 5 rpm, a gradual increase of speed up to 40 rpm and a maximum duration of 5 minutes. Fall latency was recorded. A fall was characterized by either the mouse falling directly in the underneath compartment or 2 consecutive rotations griping the rod. Mice falling from the rod due to walking backwards were excluded. The experimenter stayed more than 2 meters away while the test was running, except to get the fallen mice. Mice of different sex were run separately. The rotating rod was 3cm in diameter with ridges and mice had 6cm width of lateral space.

### Morris Water Maze

The test was conducted over 5 consecutive days. Each daily session consisted of 4 trials with the start location being semi-random between 4 cardinal locations (north, N; east, E; northwest, NW; southeast, SE): day 1- N, E, SE, NW; day 2- SE, N, NW, E; day 3- NW, SE, E, N; day 4- E, NW, N, SE; day 5- N, SE, E, NW. Location of the platform was the same for all mice, in the southwest quadrant, not close to the border nor the center of the basin. At the start of each trial, mice were gently lowered in the water manually, at the basin’s border and facing the wall. The basin was 120cm in diameter and the platform was 10 cm in diameter with grooves. The platform was underwater and not visible to the animal. The amount of water above the platform was adjusted between males and females so that mice would be able to climb on it easily. The basin water was kept clear, clean, and any debris removed prior to the start of each session.

Mice had a maximum of 2 minutes to find the platform after being introduced in the water. Upon reaching the platform by themselves or being gently led to it after the maximum 120 seconds, mice had to remain on the platform for 30 seconds. Visual cues consisting of simple geometrical shapes (square, circle, triangle, “lightning”, star, oblique stripes, diamond) of 20cm size were on the cardinal sides of the basin and other shapes another 1 to 1.5 m on the walls of the room.

The experimenter always exited the surrounding of the basin from the same point after introduction of the mouse. The experimenter was not visible to the mice and monitored the mice progress more than 2m away. After each session, excess water was removed with clean dry paper towels, and the mice placed in clean container with heating to recuperate. Mice were monitored and returned to their cage once dry and active.

### Perfusion for fresh frozen tissue

Animals were irreversibly anesthetized with an overdose of Isoflurane (Covetrus) and then an intracardiac perfusion was performed with ice-cold 7.4pH PBS – 50µg/mL Cycloheximide (Sigma).

For use of separate brain regions, the brains were dissected in ice-cold 7.4pH PBS and directly store at -80°C until further processing.

### Perfusion for immunohistochemistry

Animals were irreversibly anesthetized with an overdose of Isoflurane (Covetrus) and then an intracardiac perfusion was performed first with ice-cold 7.4pH PBS - 1mg/mL Heparin, followed by 7.4pH PBS - formaldehyde 4%. Brains were post-fixed in PBS-FORMALDEHYDE 4% overnight at 4°C before being transferred to 30% sucrose for cryoprotection and snap frozen with -45 +/- 5°C 2-methylbutarate (Isopentane). Frozen brains were then stored at -80°C until further processing. Fixed brains were sliced with a cryostat at 10µm in a sequential manner, before being put on charged slides and left to dry for at least 24h at room temperature (RT) before further use.

### Immunohistochemistry for image segmentation

After slicing, slides were left to dry at RT for at least 24h before use. Slides were rinsed in PBS for 5min at RT with shaking. Antigen retrieval was performed in10mM Citrate Buffer at pH 6.0 for 10min at 75°C. Slides were then rinse in PBS at RT for 5min. Permeabilization was performed with a 5min wash in PBS with 1% SDS (Apex Biotech) and 0.1% Triton X-100(Tx) (Sigma-Aldrich). Slides were then rinsed for 5 min 3 times in PBS + 0.1% Triton X-100. Hydrophobic contours were then done around individual sections using Liquid Blocker (Electron Microscopy Sciences). Blocking was performed for 60min at RT with PBS + 5% Normal Goat Serum + 0.3% Triton X-100). Primary antibodies (NeuN (E4M5P), Cell Signaling) were used at 1:500 dilution in PBS with 1% Bovine Serum Albumin and 0.3% Triton X-100, overnight (12-16h) at 4°C without shaking, flat, and in a moist box. Slides were then rinse in PBS for 5min, 3 times. Secondary antibodies were used at 1:500 dilution in the same dilution buffer as the primary antibodies, for 2h at RT, flat without shaking and in a moist environment. Slides were then rinsed 3 times for 5min in PBS. DAPI (Cell Signaling) was applied at 1µg/mL in PBS for 5min at RT, flat without shaking and in a moist environment. Slides were then rinsed 2 times for 5min in PBS at RT. Slides were then mounted with 90% glycerol - 10% PBS.

### Image segmentation

DAPI-NeuN stained coronal sections at Bregma -0.3/-0.7mm were selected based on their quality (incomplete or damaged sections for the area of interest were excluded). Sections were imaged with a Zeiss LSM900 microscope at 10X magnification using a wide-field setting and tiled using the associated ZEN software automatic tiling with 10% overlap. Using ImageJ, a rectangle of 3000µm x height of the neocortex located above the lateral ventricle and corresponding to primary somatosensory regions for Front Limbs and Hind Limbs (S1FL, S1HL) was selected. Medial lateral marker consisted of the disappearance of the visible layer 4 on the medial side (start of the primary motor cortex – M1). Height limits were the top and bottom of the neocortex, with very limited curvature at that location. For optimal visualization, contrast and brightness of the staining was adapted for each image. Cellpose (v2.0) was used with the pre-trained model “cyto” to locate and delineate nuclei and somatic cytoplasm. ImageJ was then used for additional measurements (coordinates, count, size). Segmentation results were visually inspected before further analysis.

### Polysome analysis

Approximately 3 mm^3^ of cortical tissue from fresh brains perfused with PBS-containing 50 µg/mL cycloheximide was homogenized in polysome lysis buffer (15 mM HEPES-KOH [pH 7.4], 7.5 mM MgCl_2_, 100 mM KCl, 2 mM dithiothreitol, 1.0% Triton X-100, 100 μg/ml cycloheximide and one tablet of EDTA-free protease inhibitors (Roche) per 10 ml) using a manual homogenizer. Lysates were normalized by protein content using Bradford reagent (Bio-Rad) and layered onto 11 ml 5–50% sucrose density gradients (15 mM Hepes-KOH, 7.5 mM MgCl_2_, 100 mM KCl, 2 mM DTT, 100 μg/ml cycloheximide, 5–50% RNase-free sucrose). Gradients were centrifuged in an SW-41Ti rotor at 36 000 rpm at 4°C for 1.5 h, and then sampled using a Biocomp Gradient Station with constant monitoring of optical density (OD) at 254 nm. A total of 1 ml fractions were collected throughout, adjusted to 0.5% sodium dodecyl sulphate (SDS) and incubated at 65°C for 5 min. One nanogram of polyA+ synthetic luciferase mRNA was added to each fraction for normalization. Samples were then treated with 200 μg/ml Proteinase K (Ambion) and digested for 45 min at 50°C, followed by 1:1 dilution with RNase-free water. RNA was extracted from diluted fractions using the hot acid phenol method, and precipitated with NaOAc and isopropanol. cDNA was prepared using the Protoscript II (NEB) reverse transcriptase with oligo dT_18_ priming according to the manufacturer’s instructions. Transcript abundance was determined by quantitative polymerase chain reaction (qPCR) using iTaq SYBR Green PCR mix (Bio-Rad) and primers specific for each transcript (see below). Measurements were then normalized to luciferase abundance and plotted as percent detected.

### Synaptosomes

Centrifugation rotor allowing only 6 tubes simultaneously, 3 animals of each group were treated together, blindly and in random order. Frozen tissue kept at -80C from similar brain regions were weighed. Tissue was then dounced in 4C 1mL pH7.4 HEPES-0.3M Sucrose buffer. First Centrifugation at 1,000g for 5min at 4C. Supernatant was carefully removed and transferred for another identical round of centrifugation. This supernatant was kept and named “total”, from which 50 µL were extracted for RNA and homogenized by pipetting with 150 µL lysis buffer from New England Biolabs Miniprep Kit (ref T2010S), other 50 µL were extracted for WB experiments and homogenized by pipetting with 150 µL of RIP buffer. RNA and WB lysate samples were kept at -80C for further processing. Remaining total lysate was then centrifuged at 17, 000g for 15min at 4C. That supernatant was carefully extracted and kept as “soma lysate” for further processing and stored at -80C. The pellet was resuspended in 4 C 1 mL HEPES-0.3M Sucrose buffer and homogenized by pipetting. This fraction was then layered gently on top of chilled 5 mL HEPES-0.8 M sucrose buffer in a centrifuge tube (Polyallomer centrifuge tubes, ref 331372 from Beckman Coulter). Tubes were balanced with the same chilled 1 mL HEPES-0.3 M sucrose buffer and centrifuged at 50,000g for 1 h at 4 C. 3mL consisting of the upper fraction and the upper part of the HEPES-0.8 M sucrose buffer were removed and discarded as containing myelin. Any present pellet was gently resuspended in the remaining buffer and layered on top of chilled 5mL HEPES-1.2 M sucrose buffer. Tubes were balanced with the same chilled HEPES-0.8 M sucrose buffer and centrifuged at 50,000g for 1 h at 4 C. The upper 3 mL were carefully collected and homogenized with chilled 5mL HEPES-0.3 M sucrose buffer and the pellet under the HEPES-1.2 M sucrose buffer was gently homogenized in HEPES-0.3 M sucrose buffer and kept as mitochondria lysate. Tubes were balanced with chilled HEPES-0.3 M sucrose buffer and centrifuged for the last time at 20,000g for 30min at 4C. This last pellet was resuspended in 200 µL HEPES-0.3 M sucrose buffer, with 100 µL used for RNA and gently homogenized with 200 µL lysis buffer from New England Biolabs Miniprep Kit (ref T2010S), while the remaining 100 µL were gently homogenized with chilled RIPA buffer. Both synaptosomes lysate samples were stored immediately at -80C for further processing.

### RNA analysis by quantitative PCR (qPCR)

RNA was extracted using the Monarch Total RNA Miniprep Kit from New England Biolabs (ref T2010S) following the part 2 without the optional DNase 1 optional treatment step. Reverse transcription was performed with the following reagents: 12µL RNA, 1µL Firefly Luciferase RNA (1pg), 1µL of 50µM oligo(dT)18 were mixed and incubated at 65C for 5min. Each tube was then complemented with the mix of the following reagents: 4µL of 5X Protoscript II buffer, 1µL of 10mM dNTPs, 1µL of 0.1M DTT, 1µ of Protoscript II RT (all reagents from New England Biolabs). The tubes were then incubated for 1h at 42C, followed by 20min at 65C and finally, 79µL of RNase-free water was added to each tube. The samples were kept at -20C for further use. Three technical replicates were made for the qPCR reactions. Wells of a 96-well qPCR plate were used with 5µL of iTaq Universal SYBR Green Supermix, 0.6µL of 5µM forward and reverse qPCR primers, µL of RNase-free water and 2.5µL cDNA. Well solutions were previously carefully homogenized, and the plate centrifuged briefly to avoid the presence of bubbles. The qPCR was conducted with an initial step of 2min at 50C, followed by an initial denaturation of 2min at 95C, then 40 cycles of: 15s at 95C, 20s at 58C and 30s at 72C; with a final step of 30s more at 72C. Cq values were quantified using the real-time PCR system (Stratagene Mx3005P from Agilent Technologies with the software MxPro). Raw Cq values were kept if the SD of the triplicates was below 0.3. Genes of interest were subtracted to the raw value for the gene of reference. The results were then normalized to either the WT total lysate average values (total and synaptosomes / total ratios) or the WT synaptosome lysate values (synaptosomes). Results were then transformed to linear scale by raising 2 to the negative normalized value.

**Table 2.**
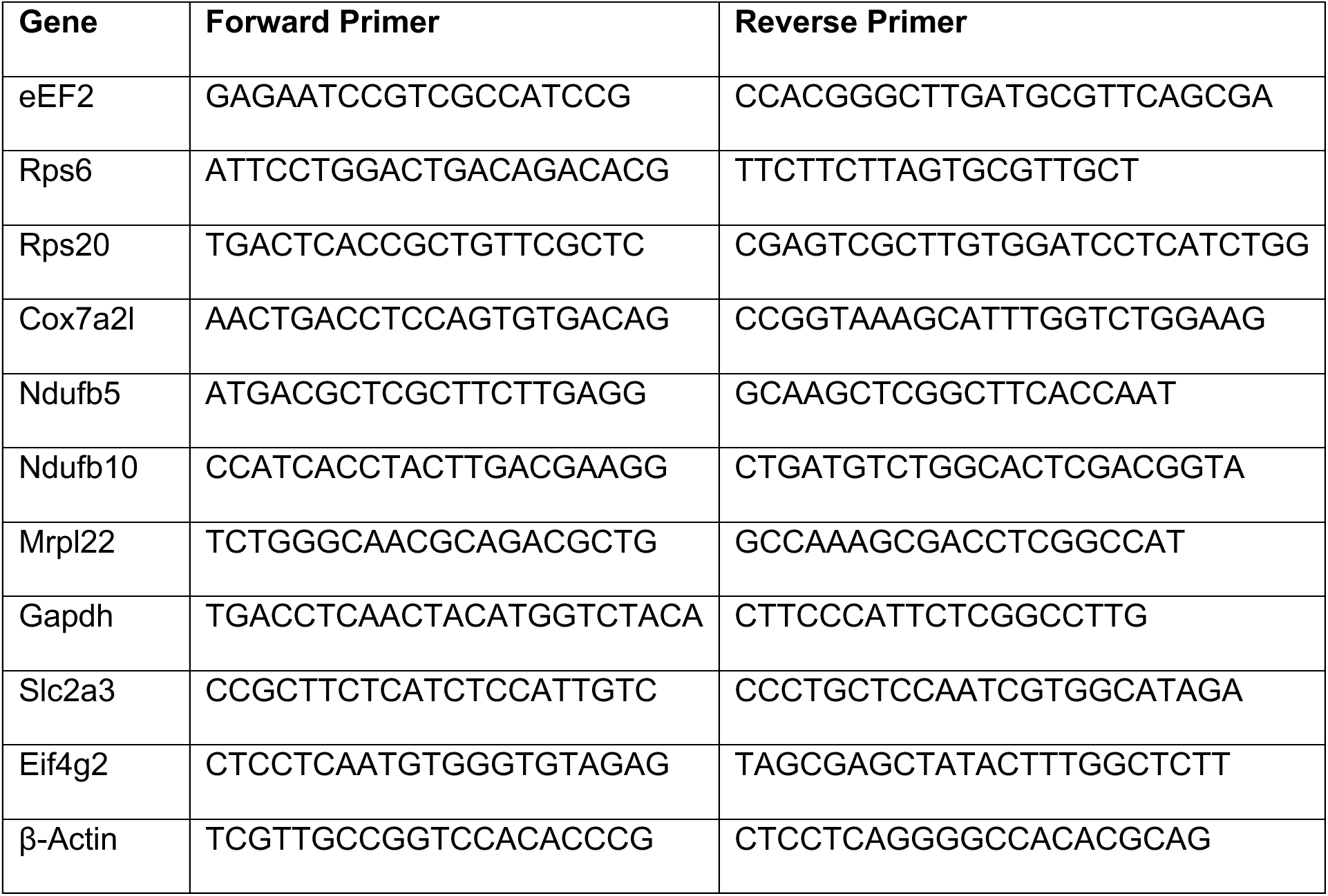
Sequences of qPCR primers for mouse mRNAs.

### RNA-seq library preparation and analysis

Total RNA was isolated from homogenized posterior cortex from 3 male WT and 3 male Larp1 cKO mice using RNEasy columns (Qiagen). rRNA was removed using the NEBNext rRNA removal v2 kit (NEB E7400) and paired-end RNA-seq libraries were prepared using the NEBNext Ultra II RNA library prep kit (NEB E7770) with Illumina primers. Synaptosome RNA-seq libraries were prepared at the Yale Center for Genome Analysis from RNA isolated from synaptosomes, as described above. All samples were sequenced on an Illumina NovaSeq X Plus at the Yale Center for Genome Analysis. Adapter sequences were trimmed using cutadapt (https://github.com/marcelm/cutadapt/), after which sequences were aligned to the mm10 genome using STAR aligner (https://github.com/alexdobin/STAR). Gene-level quantification was performed use RSEM (https://github.com/deweylab/RSEM) with UCSC gene annotations, and differential expression was quantified using the DESeq2 software package (59). Coding mRNAs were identified according to GENCODE VM36 annotations. RNA-seq libraries for WT and LARP1 KO HEK-293T cells were sequenced previously (4), aligned to hg38 using the STAR aligner, and quantified using GENCODE43 gene annotations using RSEM. Differential expression between WT and KO HEK-293T cells was calculated using DESeq2, as above.

### Analysis of transcription start sites and calculate of TOPscores

Transcription start sites were extracted from HeliScopeCAGE (hCAGE) data downloaded from the Fantom5 project (http://fantom.gsc.riken.jp/5/). Transcription start sites in the mouse brain were determined using hCAGE results in <u>cortex%2c%20neonate%20N30.CNhs11107.1392-42F2.mm9.nobarcode.bam</u>. TOPscores for mouse brain were determined using this file and custom scripts described previously (4). TOPscores for HEK-293 cells were calculated previously (4).

### Western Blotting

Samples were mixed with 5X protein denaturing buffer and incubated at 98C for 2min. Denatured samples (15 µL) were then loaded on to a NuPAGE 4-12% Bis-Tris Gel from Invitrogen (NP0323) and run with SDS MOPS at 130 C approximatively 100 min or until the lower ladder band reached the bottom. Gel was then transferred to a membrane by running it for 2h at 45 V in homemade transfer buffer. Membranes were then blocked in 5% milk-TBST for 1 h at RT, before being rinsed in TBST for 5min. Primary antibodies were diluted as indicated in 5% BSA-TBST and the membranes incubated with gently rocking at 4C overnight. Membranes were then washed 3 times for 5 min each with rocking. They were then incubated with a secondary antibody for 1 h at RT. Membranes were then washed 3 times for 5 min each. Membranes incubated with a fluorescent secondary antibody were then imaged on a LI-COR Odyssey and quantified with Image Studio 6. Membranes incubated with an HRP secondary antibody were then incubated with Clarity Western ECL substrates (#170-5061 from Bio-Rad) for 5 min and imaged on film.

## Acknowledgements

We thank all members of the Thoreen lab for discussion and critical reading of the manuscript. We also thank Susumu Tomita and Marcelo Dietrich for reagents, discussions and critical feedback. We are additionally grateful to Ken Inoki for providing the Larp1^flox/flox^ mouse. This work was supported by the U.S. National Institutes of Health (R35GM152167 to C.C.T and R21NS118616 to C.C.T, http://nih.gov). The sponsors played no role in the study design, data collection and analysis, decision to publish or preparation of this manuscript.

## Author Contributions

M.S.W and C.C.T conceived the project. M.S.W performed experiments. M.J.W conducted polysome and RNA-seq analysis. C.C.T supervised the study. All authors discussed the results and edited the manuscript.

## Data Availability

Raw and analyzed data from all high-throughput sequencing experiments have been deposited on the Gene Expression Omnibus (https://www.ncbi.nlm.nih.gov/geo/) under accession number GSE286175.

## Supplemental Materials

**Supplemental Table 1.** Differential gene expression between Larp1 WT and cKO cortex.

**Supplemental Table 2.** TOPscores for genes in mouse cortex.

**Supplemental Fig 1.**
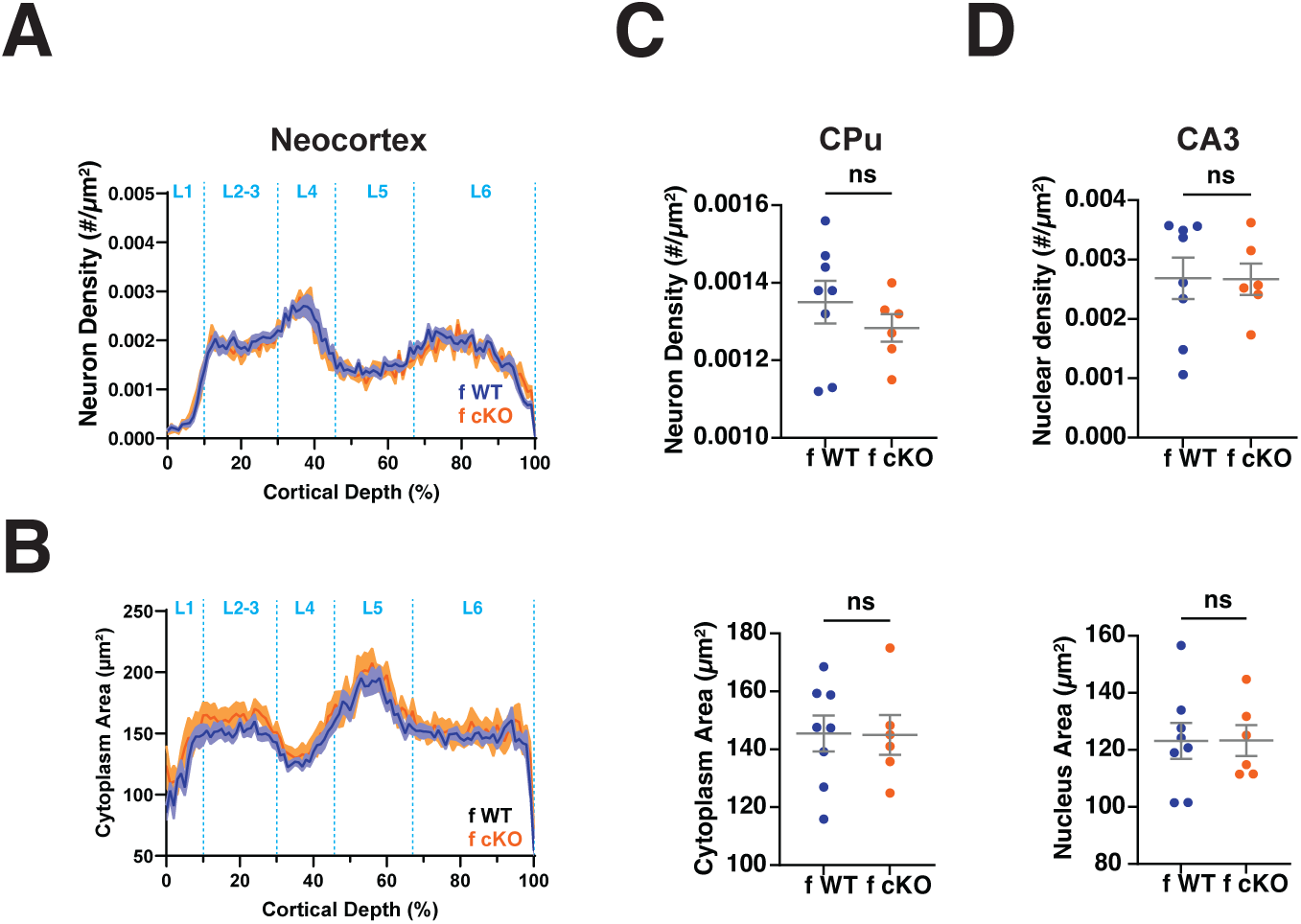
Neuron size and density in female mice. **A.** The density of cortical neurons in Larp1 WT and cKO female mice. NeuN+ cells from regions of the primary somatosensory region from female Larp1 WT and cKO mice were segmented and binned by position along the vertical axis in 1% increments. Plot shows mean number and SEM within each bin. Significance by REML group, n=8 WT and 6 cKO. **B.** Sizes of neuronal soma in female Larp1 WT and cKO mice. Areas of NeuN+ cells in segmented regions from (A) for female Larp1 WT and cKO mice. Traces are means +/- SEM. Significance by REML group p value = 0.0257, n=8 WT and 6 cKO. **C.** Neuronal density in the caudate putamen of female Larp1 WT and cKO mice. NeuN+ cells in the caudate putamen (CPu) of female Larp1 WT and cKO mice were segmented and analyzed as in (A). Top panel: density of NeuN+ cells from the indicated genotypes. Significance by t-test, n n=8 WT and 6 cKO. Bottom panel: area of NeuN+ cells from the indicated genotypes. Significance by t-test, n=8 WT and 6 cKO. **D.** Density of nuclei in the CA3 region of the hippocampus of female Larp1 WT and cKO mice. DAPI+ nuclei from the CA3 region were segmented and analyzed as in (A). Top panel: density of DAPI+ nuclei in the indicated genotypes. Significance by t-test, n=8 WT and 6 cKO. Bottom panel: area of DAPI+ nuclei from the indicated genotypes. Significance by t-test, n=8 WT and 6 cKO.

**Supplemental Fig 2.**
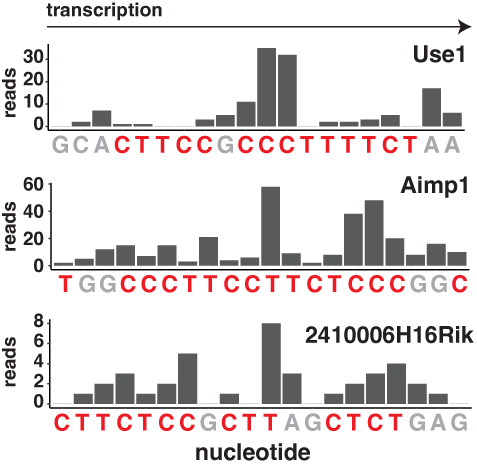
Transcription start sites of non-canonical mRNAs. Examples of TOP motifs in mRNAs depleted from Larp1 cKO mice. Transcription start sites for the indicated genes from FANTOM5 CAGE data analysis of mouse neonatal cortex (see Methods).

**Supplemental Fig 3.**
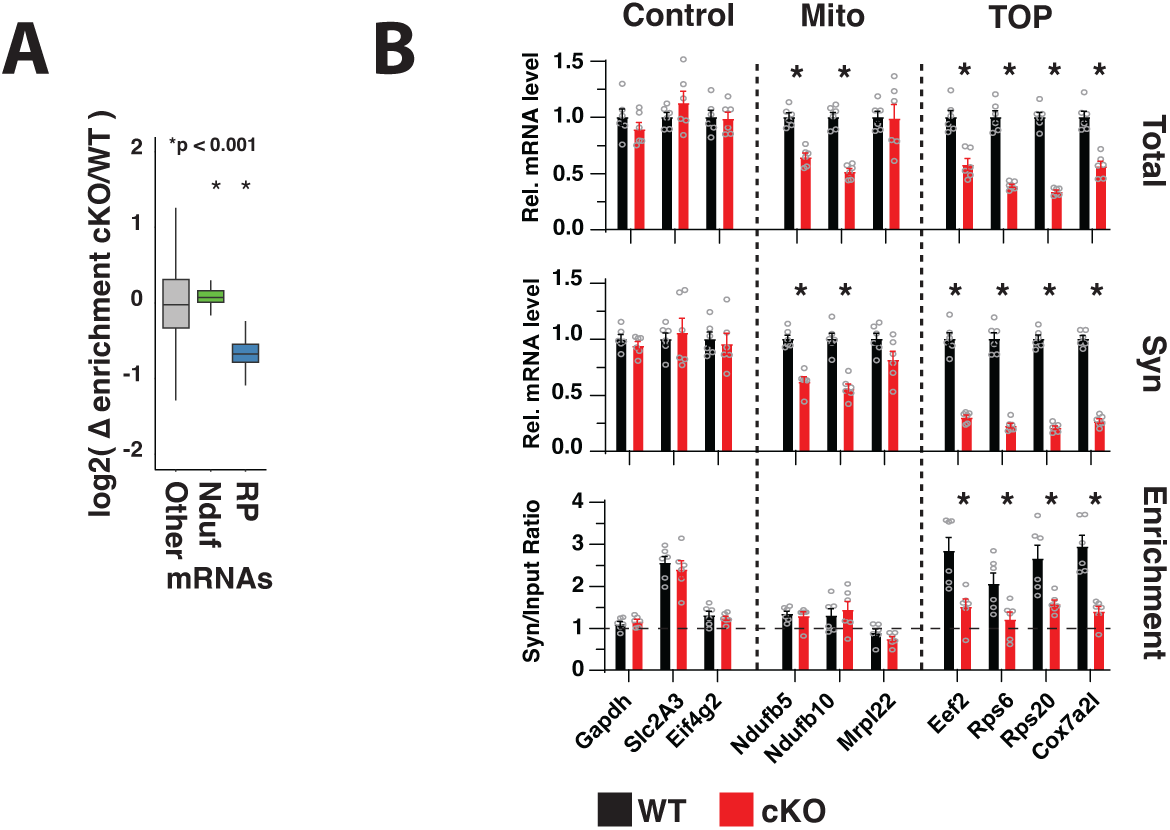
Synaptosome enrichment of TOP and non-TOP mRNAs in synaptosomes from WT and Larp1 cKO brains. **A.** Complex I (Nduf) mRNAs are not depleted from synaptosomes in Larp1 cKO brains. Plot shows changes in the synaptosome enrichment (synaptosome/total) of Nduf, ribosomal protein (RP) and other mRNAs between WT and Larp1 cKO brains. Significance by t-test comparison between each class and Other mRNAs. **B**. Validation of changes in synaptosome enrichment in Larp1 cKO brains by qPCR. Levels of the indicated mRNAs in input and synaptosome samples from Larp1 WT and cKO brains were quantified by qPCR (n=6 for each genotype, error bars are SEM, significance by unpaired t-tests).

